# Automated classification of giant virus genomes using a random forest model built on trademark protein families

**DOI:** 10.1101/2023.11.10.566645

**Authors:** Anh D. Ha, Frank O. Aylward

## Abstract

Viruses of the phylum *Nucleocytoviricota*, often referred to as “giant viruses,” are prevalent in various environments around the globe and play significant roles in shaping eukaryotic diversity and activities in global ecosystems. Given the extensive phylogenetic diversity within this viral group and the highly complex composition of their genomes, taxonomic classification of giant viruses, particularly incomplete metagenome-assembled genomes (MAGs) can present a considerable challenge. Here we developed TIGTOG (<u>T</u>axonomic Information of <u>G</u>iant viruses using <u>T</u>rademark <u>O</u>rthologous <u>G</u>roups), a machine learning-based approach to predict the taxonomic classification of novel giant virus MAGs based on profiles of protein family content. We applied a random forest algorithm to a training set of 1,531 quality-checked, phylogenetically diverse *Nucleocytoviricota* genomes using pre-selected sets of giant virus orthologous groups (GVOGs). The classification models were predictive of viral taxonomic assignments with a cross-validation accuracy of 99.6% to the order level and 97.3% to the family level. We found that no individual GVOGs or genome features significantly influenced the algorithm’s performance or the models’ predictions, indicating that classification predictions were based on a comprehensive genomic signature, which reduced the necessity of a fixed set of marker genes for taxonomic assigning purposes. Our classification models were validated with an independent test set of 823 giant virus genomes with varied genomic completeness and taxonomy and demonstrated an accuracy of 98.6% and 95.9% to the order and family level, respectively. Our results indicate that protein family profiles can be used to accurately classify large DNA viruses at different taxonomic levels and provide a fast and accurate method for the classification of giant viruses. This approach could easily be adapted to other viral groups.

## Introduction

Large viruses of the phylum *Nucleocytoviricota*, commonly referred to as “giant viruses”, are a diverse group of double-stranded DNA eukaryotic viruses with large virion sizes, reaching dimensions of up to 1.5 μm, which is comparable to the sizes of several archaea, bacteria, and eukaryotes^1–4^. These viruses have been discovered ubiquitously in the biosphere and potentially play key roles in shaping the structure of microbial communities and biogeochemical cycling^5–10^. Currently, documented members of the phylum can be divided into five orders: *Algalvirales, Asfuvirales, Chitovirales, Imitevirales,* and *Pimascovirales,* as well as 11 established and potentially many new potential families^11,12^. Nucleocytoviruses are known to infect a wide range of eukaryotic hosts; whereas members of the *Algavirales* and *Imitervirales* orders infect diverse algae, amoebae and other protists, members of the *Asfuvirales, Chitovirales,* and *Pimascovirales* infect a mixture of metazoan and protist hosts^3,13–16^. Their genome sizes encompass an exceptionally wide spectrum, ranging from less than 100 kbp to over 2.7 Mbp^17,18^. Previous comparative genomic analyses have highlighted the exceptional complexity of giant virus genomes and suggested dynamic gene exchanges between these viruses and their host cells, as well as with other viruses^9,19–21^. Within the *Nucleocytoviricota* phylum, substantial phylogenetic diversity among members has been observed, and recent metagenome-enabled studies have vastly expanded the known diversity of this group^22,23^. Due to the remarkably large phylogenetic breadth and the chimeric nature of their genomes, taxonomic classification of giant viruses presents a considerable challenge.

To date, phylogenetic analyses of giant viruses have primarily relied on analysis of a small set of core genes^24,25^. While alignment-based approaches have proven effective, they present challenges in numerous instances^26^. Most notably, construction of large phylogenetic trees requires multiple computationally-intensive steps, including multi-sequence alignment and tree inference. Manual analysis of large trees to taxonomically assign a few genomes is oftentimes impractical due to the substantial time and computational resources required. Furthermore, it is not uncommon for novel genomes assembled from metagenomes to be incomplete. The process of reconstructing giant virus genomes from metagenomic reads, including *de novo* assembly and binning steps, is prone to errors. As a result, it can potentially produce fragmented MAGs that lack predicted proteins with matches to the traditional marker gene set, thereby compromising the ability to produce good-quality alignments. Due to these challenges, methods that do not require the construction of phylogenetic trees have emerged as promising alternatives^27–29^. Many of these approaches also make use of machine learning, which has been shown to be successful in a range of applications, including virus identification and classification^29–38^. Machine learning algorithms can be considerably less computationally demanding compared to methods that require multi-sequence alignment and tree inference, allowing the trained models to be effectively applied to large query datasets that would otherwise be impractical to handle^39^.

Here we developed TIGTOG (<u>T</u>axonomic Information of <u>G</u>iant viruses using <u>T</u>rademark <u>O</u>rthologous <u>G</u>roups), a machine learning-based approach to classify novel giant virus genomes based on a broad genomic signature, rather than relying on a fixed set of marker genes. We trained TIGTOG with a diverse genome dataset that consisted of sequences from all major documented phylogenetic lineages of giant viruses and other large viral groups often found to bear similarities to giant viruses. We tested our classification model using an independent test set of viral genomes with varying levels of genomic completeness and taxonomy. Our work provides a rapid, reliable tool to identify the taxonomic assignments of novel giant virus genomes and potentially other viral groups.

## Results and Discussion

### Construction of classification models

TIGTOG employs a machine learning approach based on protein family profiles to classify giant virus genomes at the order and family level. Genomes within the *Nucleoviricota* phylum exhibit high diversity and distinct signatures of protein content among different taxonomic groups (Fig. 1, Fig. S1). Each lineage harbors a unique set of protein families, i.e., distinct giant virus orthologous groups (GVOG) composition, that can be leveraged for classification purposes. We hypothesized that these unique protein family profiles could provide predictive information for the taxonomic classification of a novel genome. Aiming to search for a classification method that relies less on a fixed set of marker genes, we employed a supervised machine learning approach using the Random Forest (RF) algorithm. The presence of ortholog groups and the GC content of sequences in the training set were used as features, and pre-established taxonomic information was used as labels during the model construction process. We chose not to use genome size as a feature because many metagenome-derived genomes can be incomplete or harbor redundancies (i.e., multiple closely related viruses binned together), and we therefore wanted to exclude these possible biases from our classification method. We included a total of 1,531 genomes belonging to all established families of the *Nucleoviricota*, *Mirusviricota*, and large *Caudovirales* (jumbo phages) in the training set. The recently-discovered *Mirusviricota* lineage is a widespread group of large DNA viruses with a herpesvirus-like capsid that represent a lineage distinct from the *Nucleocytoviricota*. Their genomes appear to contain elements from various viral lineages, including the *Nucleocytoviricota*, and may therefore be misclassified as giant viruses based on the genomic contents. Jumbo phages are tailed bacteriophages with genomes exceeding 200 kbp in size and can have a misleadingly high number of giant virus orthologous group matches^40,41^. Viruses of the *Mirusviricota* and jumbo phages were therefore incorporated into our training databases because they are a likely source of false-positive classification as giant viruses.

**Figure 1.**
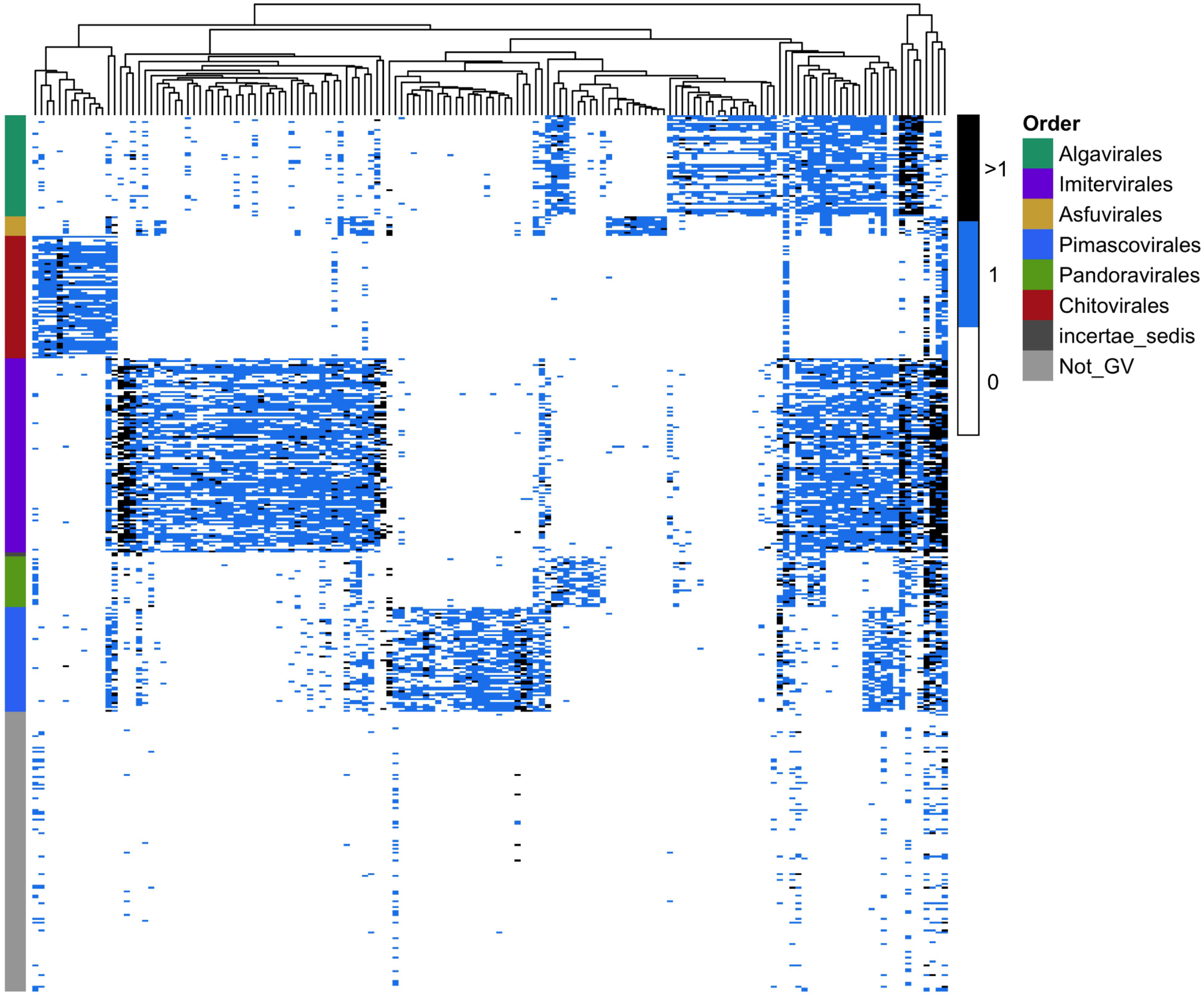
Distinct protein family profiles in different *Nucleocytoviricota* orders. The y-axis denotes the taxonomic order of giant virus representative genomes, color-coded by their respective order. The x-axis shows different GVOGs, which were used as features during the training of the classification model at the order level. The Not_GV group includes *Mirusviricota* and jumbo phage genomes.

The workflow for the machine learning pipeline is described in Fig. 2. For the first round of training, we identified a set of 625 GVOGs that are found in at least 25% of the genomes in each order. We built initial RF classification models on the order and family levels, using all 625 GVOGs and the sequences’ GC content as features. We tuned the models using randomized search cross-validation followed by grid search cross-validation. Optimal hyperparameters for models were selected through a 10-fold grid search cross-validation. We arrived at two initial models with classification accuracy of 99.7% at the order level and 97.6% at the family level.

**Figure 2.**
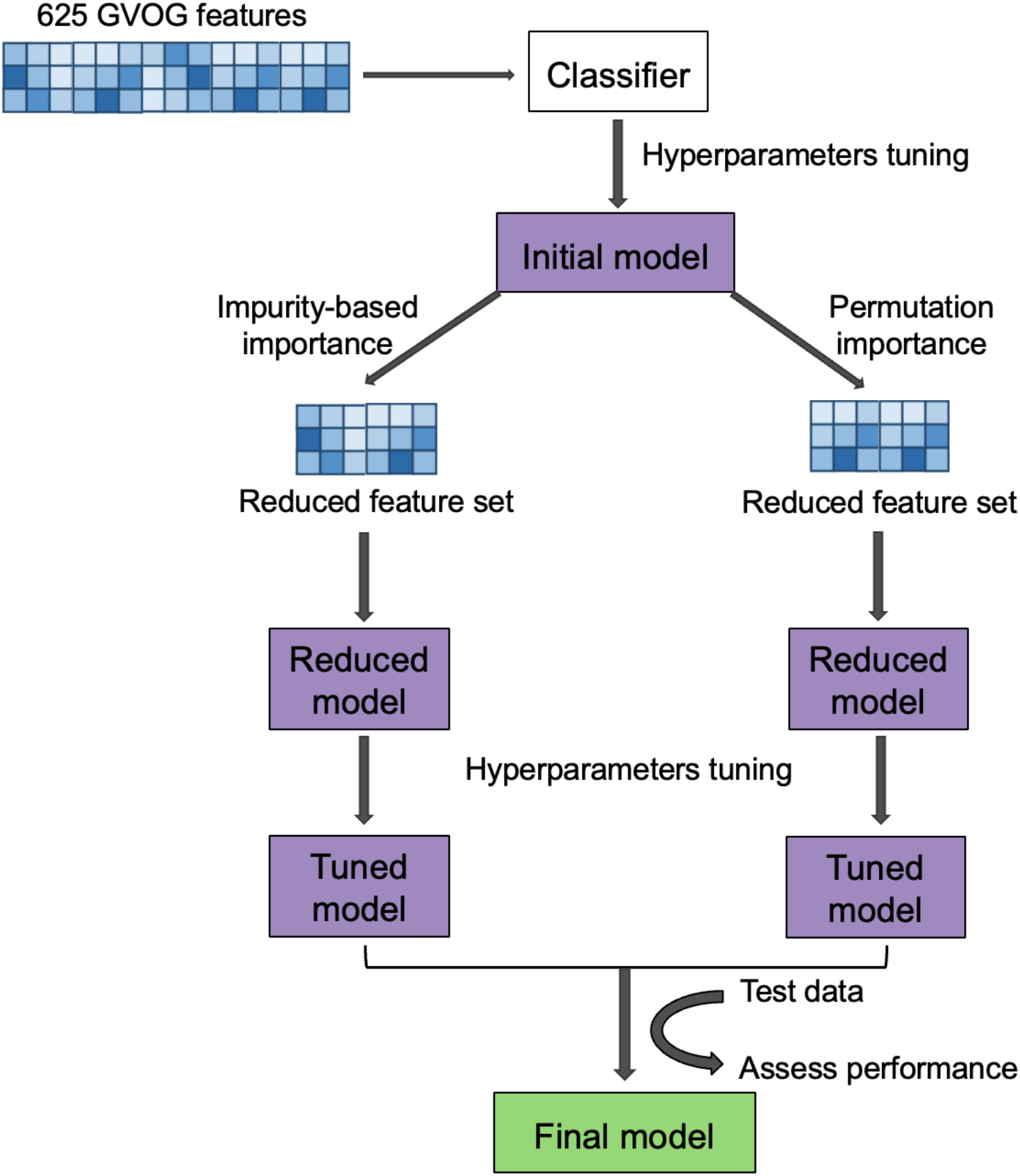
Overview of the model training pipeline at each taxonomic level. (order and family)

To further probe the characteristics of these models, we examined whether the prediction accuracy was affected by the number of GVOG features employed, using Recursive Feature Elimination (RFE) with cross-validation. We observed a plateau of accuracy scores, as indicated by similar mean values and overlapping error bars from approximately 150 and 200 features onwards for the order- and family-level classifiers, respectively (Fig. 3A). Expanding the feature set beyond these values did not yield a significant improvement in the model’s accuracy. This may be attributed to the presence of correlated relationships among the GVOG features. As correlated features can provide redundant and similar information, the inclusion of many repetitive, non-informative features does not contribute significantly to the classifiers’ performance, and may even lead to over-fitting. Indeed, hierarchical clustering based on the Spearman rank-order correlations indicated strong collinearity within the GVOG data matrix (Fig. S2). This implies that the optimal number of features for the model might fall within this range, negating the need to include the entire set of 625 GVOGs.

**Figure 3.**
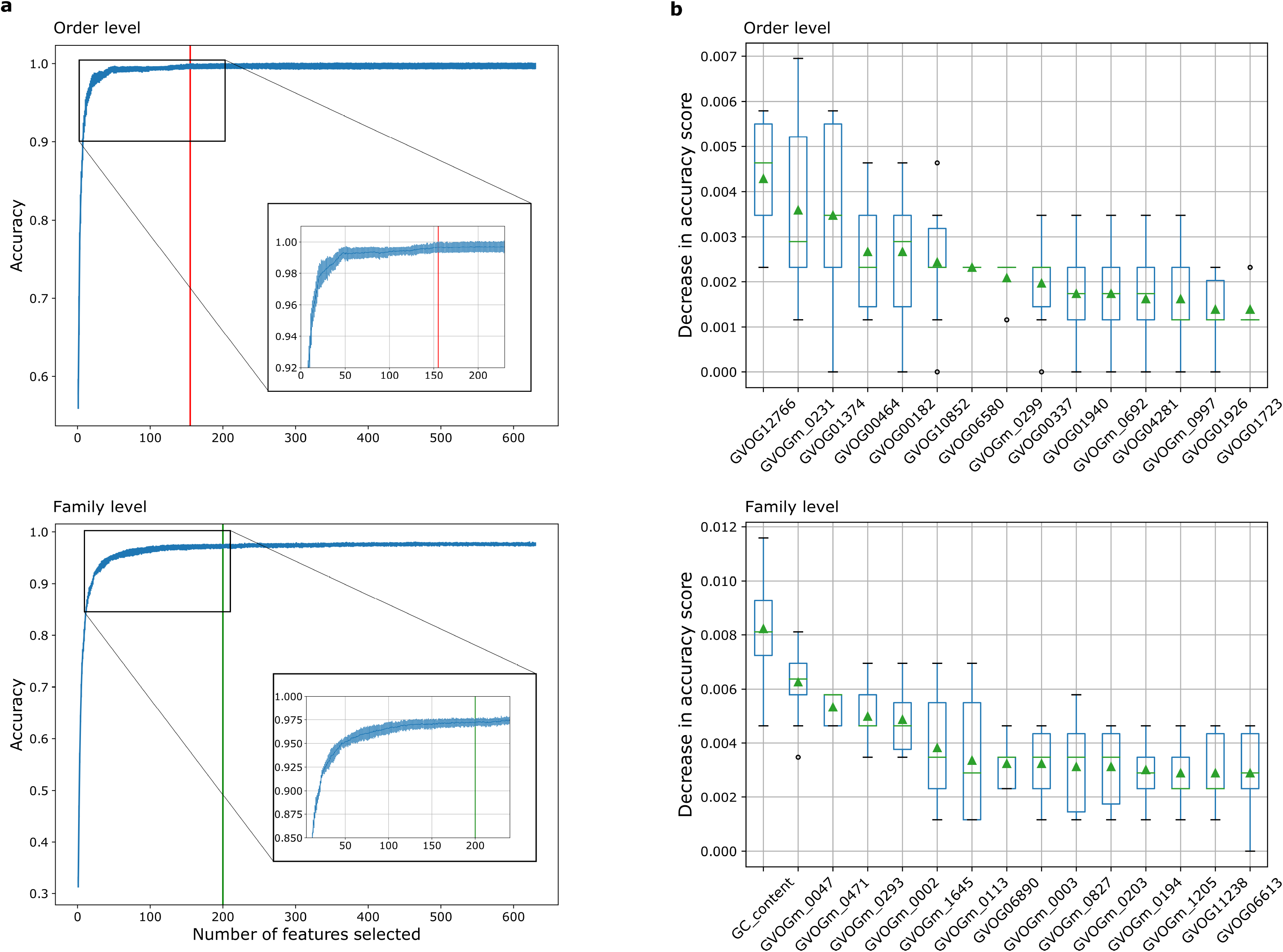
Evaluation of the impact of the initial 625 GVOG feature set on the random forest algorithm for predicting taxonomy. (a) Changes in the prediction accuracy scores with increasing number of features at the order level (top) and the family level (bottom). The vertical lines indicate the number of GVOGs that were employed in TIGTOG’s final models. (**b)** Permutation importance for the 15 most important features in the classification model at the order level (top) and family level (bottom). Features were shown in decreasing order based on their impact on accuracy when they were randomly permuted. Permutation importance testing was performed 10 times. Mean values were denoted by green triangles.

Conducting HMM searches against a large set of HMM profiles is a computationally demanding and time-consuming task. To mitigate the computational expense during the data preparation step, we aimed to reduce the GVOG set and select only the optimal features needed for the models’ performance. We ranked the feature importance using permutation importances, which measures the mean decrease in model’s prediction accuracy when a feature is randomly permuted. In general, individual features showed low importance in the model’s performance; permutation of the most important GVOG feature caused only a marginal average decrease of 0.4% at the order level and 0.6% at the family level in accuracy scores (Fig. 3B). This suggests that no single GVOG has a particularly strong influence on the model’s prediction. To further confirm this observation, we applied an alternative measure of feature importances based on the mean decrease in impurity (MDI). Impurity-based feature importances can potentially be misleading, especially when applied to predictor variables with varying measurement scales and numerous categories^42^. Nevertheless, in specific situations, including highly correlated data, RF variable importance measures could still provide valuable insights^43,44^. MDI also suggested the same result as individual features showed low importance in the model’s performance (Fig. S3). These results suggest that effective taxonomic classification could be based on broad genomic signatures, which lessens the necessity of a fixed set of marker genes for taxonomic assigning purposes.

The features with the highest importance scores identified by each of the above feature selection methods were subsequently extracted from the 625-feature set and passed to a new RF model for training. We estimated the performance of the models using 10-fold nested cross-validation, which provided an estimation of each model’s ability to generalize to unseen data. We chose the models with feature sets selected through impurity-based importance as our final classifiers for TIGTOG, as they demonstrated better performance. While the models trained with feature set selected based on MDI yielded average nested cross-validation accuracies of 99.6% at the order level and 97.3% at the family level, the models based on permutation importance had mean accuracies of 98.1% and 96.1%, at the order and family level, respectively (Fig. S4).

The feature sets selected for the final models included marker genes that are prevalent across all giant virus groups, but absent in non-*Nucleocytoviricota* genomes, such as GVOGm0003 (giant virus major capsid protein), GVOGm0760 (packaging ATPase), GVOGm0890 (Poxvirus late transcription factor VLTF3), GVOGm0032 (Ser/Thr protein phosphatases), GVOGm0095 (D5-like primase), and other more lineage-specific genes (Fig. 1, Supplementary Data S1).

Next, we assessed whether the performance of the final TIGTOG models was influenced by the number of sequences included in training. We examined how the order- and family-level final models’ cross-validation accuracy changed with an increasing training set size (Fig. 4). Generally, the performance of the models improved as the number of training instances increased. At the largest number of training sequences, cross-validation accuracy reached 99.6% at the order level and 98.2% at the family level. The learning curves suggested that adding more training examples was likely to improve models’ cross-validation accuracy at both taxonomic levels. It is possible that the unequal representation of taxonomic groups within the training dataset may contribute to this trend. For example, at the order level, there were only 102 *Chitovirales*, 66 *Asfuvirales*, and 18 *incertae sedis* sequence instances in the training data, in contrast to the most abundant group *Imitervirales*, which included 2,782 sequences. At the family level, many families were represented by 30 or fewer sequences, while the family *Imitervirales 01* had 1,788 representatives. Adding more sequences from under-represented groups could potentially provide more information about these groups’ genomic signature and account for more diversity within the group, therefore improving the overall accuracy of the models’ predictions.

**Figure 4.**
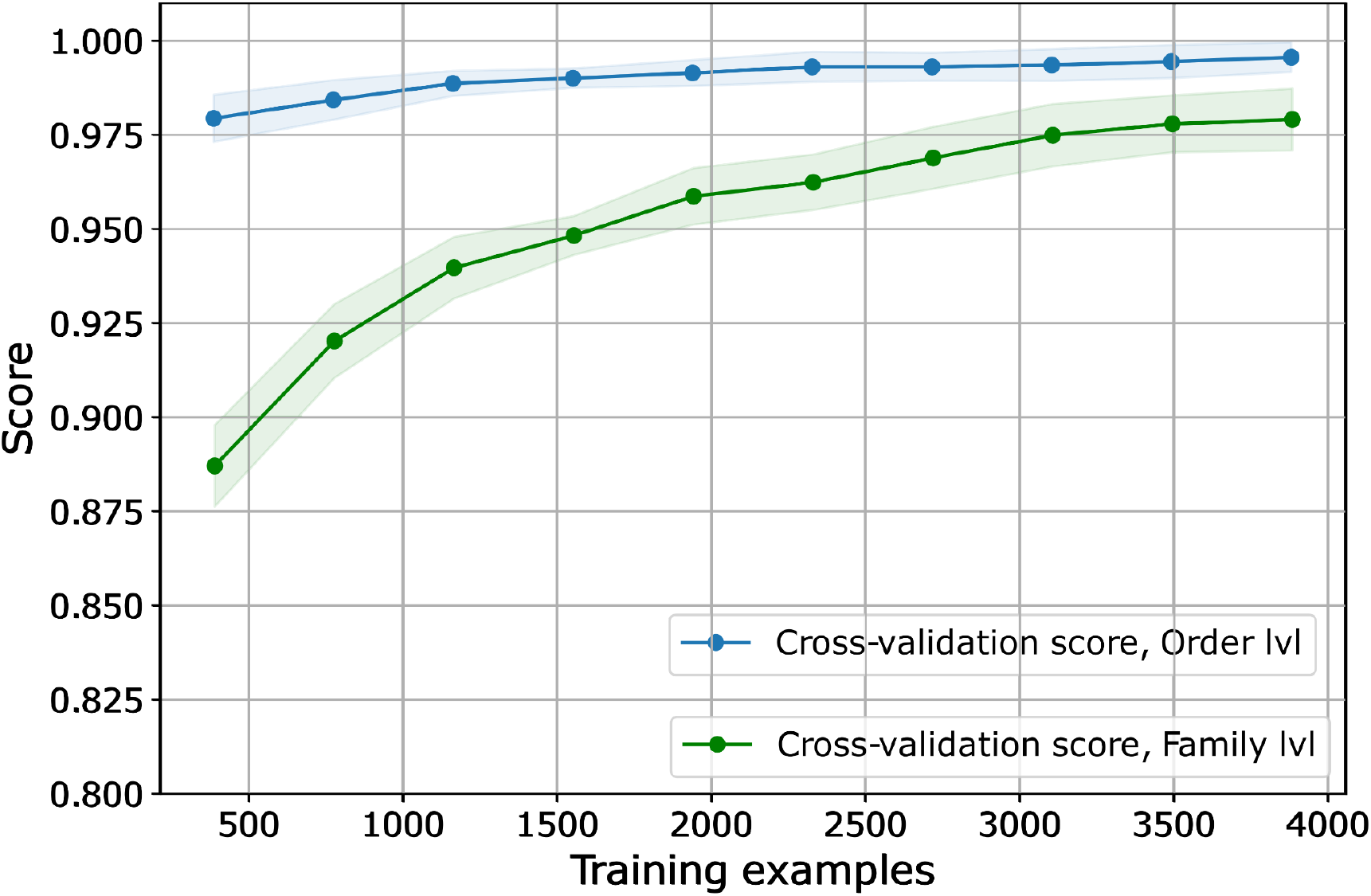
The performance of final classification models over varying number of training instances. The curves are plotted with the mean cross validated test scores. Shaded areas represent a standard deviation above and below the mean for all cross-validations. The scores of models at the order level is in shown blue and scores of model at the family level is in green.

### Evaluating models’ predicting taxonomic classification based on sequence content

We next assessed the models constructed using the training genome set to predict the taxonomic classification of each genome in the independent test set. In this test set, we included representatives from all *Nucleoviricota* families, along with genomes from the *Mirusviricota*, jumbo phages, and virophages as examples of non-giant-virus sequences. Additionally, we introduced fragmented sequences at various completeness levels, as described in the Methods section. The models demonstrated a sufficient ability to generalize to new data (Fig. 5A). The performance of the model on the test set at the order level was comparable to the estimates made through nested cross-validation, with an accuracy of 98.6%. At the family level, the model’s prediction achieved an accuracy of 95.9%. This suggests that TIGTOG are broadly applicable when tested against diverse sequence groups.

**Figure 5.**
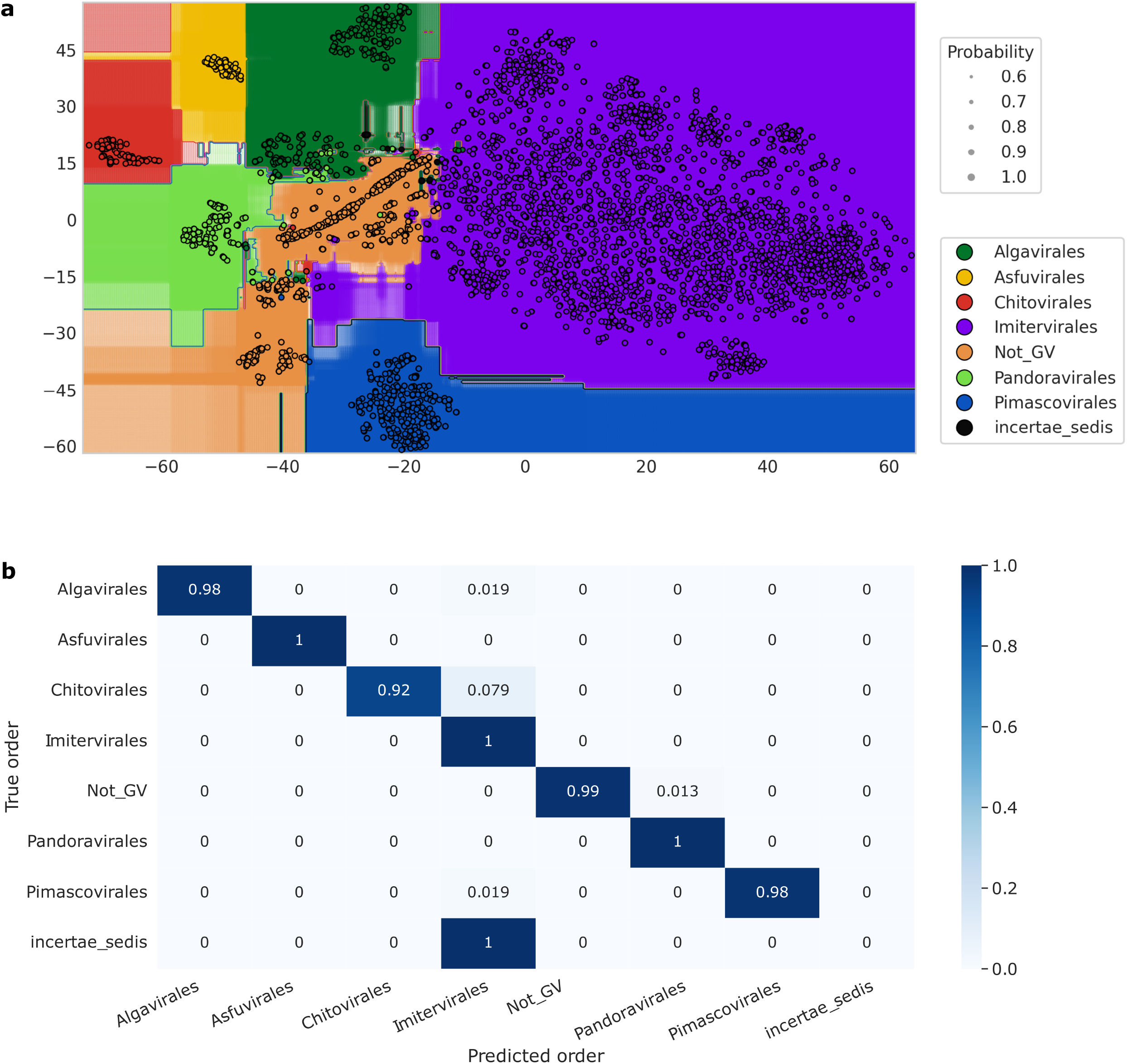
Evaluation of the final classification model’s performance at the order level. (a) Decision boundary plotted for classifier at the order level in dimension of two t-distributed stochastic neighbour embedding (T-SNE) components. Data dimensions were reduced using PCA and T-SNE. All dots are colored by the giant virus order. Training data are visualized in circles with black border. The sizes of the transparent dots (without border) indicate probability of class membership for each point on the grid across the feature space. **(b)** Normalized confusion matrix of classification at the order level. Rows correspond to the true taxonomic assignments of sequences, and columns represent predicted classification. The diagonal values indicate the percentage of times the predicted classification matches the true taxonomy. Values were normalized by class sizes.

At the order level, out of 823 test genomes with varied levels of completeness, 11 genomes (1.3%) were classified incorrectly (Fig. 5B). Four out of these 11 sequences were simulated incomplete genomes derived from the other seven MAGs. Among these, seven sequences had a completeness level of less than 70%. The sequences that were falsely classified had completeness levels ranging from 45% to 100% compared to the original sequences; this indicates that the accuracy of TIGTOG was not significantly affected by the completeness of the sequences. In 9 out of 11 incorrect instances, TIGTOG misclassified sequences as *Imitervirales.* Table 1 details a classification report of the model’s prediction at the order level. In general, the classifier performed adequately (F1 score >= 0.96 for all classes), with the exception of the *incertae sedis* genomes, where two sequences included were both incorrectly classified into *Imitervirales*. This suggests a potential issue with class imbalance in the training dataset, where the model may exhibit bias towards *Imitervirales,* the most populous class.

**Table 1.** Classification report for model’s prediction at the order level. The weighted-averaged scores were calculated by taking the mean of all per-class scores while considering the support for each class.

|  | Precision | Recall | F1-score | Support |
| --- | --- | --- | --- | --- |
| *Algavirales* | 1 | 0.98 | 0.99 | 52 |
| *Asfuvirales* | 1 | 1 | 1 | 10 |
| *Chitovirales* | 1 | 0.92 | 0.96 | 63 |
| *Imitervirales* | 0.98 | 1 | 0.99 | 463 |
| Not_GV | 1 | 0.99 | 0.99 | 153 |
| *Pandoravirales* | 0.93 | 1 | 0.96 | 26 |
| *Pimascovirales* | 1 | 0.98 | 0.99 | 54 |
| *incertae_sedis* | 0 | 0 | 0 | 2 |
| **Accuracy** |  |  | 0.99 | 823 |
| **Weighted avg** | 0.98 | 0.99 | 0.99 | 823 |

At the family level, 34 out of 823 test genomes (4.1%) were incorrectly classified. 13 out of these 34 sequences were simulated incomplete MAGs derived from the other 21 genomes. All these incorrect predictions displayed relatively low confidence, with confidence levels ranging from 12% to 63%, and the majority (33 out of 36 incorrect predictions) had a confidence level below 50%. Hence, the reported confidence level may serve as a good reference, particularly in the prediction of families. The family-level classifier only recognizes potential members of the major families, such as AF_01, AG_01, IM_01, IM_16, PM_01, PV_04, and PX_01 (details of all the major families that TIGTOG recognizes are listed in Fig. S1). Other families with fewer representatives were combined into a group labeled with the order name (e.g., AF, AG, IM, PM, and PV). As genomic content can vary significantly between families, this aggregation unsurprisingly resulted in lower precision and recall scores for the groups without family notations (Supplementary Data S3).

### Comparison of input sequences to reference giant virus database using average amino acid identity (AAI) calculation

In addition to taxonomic assignment, TIGTOG can also perform protein similarity calculations between input sequences and established giant virus genomes using LAST searches. The custom reference database included a wide phylogenetic variety, containing representatives of every *Nucleocytoviricota* genus as previously described (details of taxonomy available in Supplementary Data S4).

## Conclusion

Overall, our results provide evidence that application of a random forest model to protein family profiles can effectively classify novel giant virus genome sequences. Our application of this approach relies on reduced-scale HMM searches against pre-selected GVOG databases, which are time-efficient and capable of handling a large number of sequences. Given the widespread distribution of giant viruses in the environment and the continuous generation of new sequences through metagenomic data, the number of newly identified giant virus MAGs is growing exponentially. An efficient classification tool would benefit ongoing efforts to characterize the environmental diversity, explore the geographic and temporal variability of these viruses in global ecosystems, and to gain deeper insights into the evolutionary traits within this phylum. TIGTOG is capable of working with incomplete sequences, and so we anticipate that this tool will be broadly useful for analyzing giant virus diversity in the biosphere. Moreover, our results provide a useful proof-of-concept that this approach can be useful for classification of other large DNA viruses.

## Materials and methods

### Genome database compilation

We compiled a database of 1,382 *Nucleocytoviricota* genomes from the Giant Virus Database (GVDB)^11^ and 696 large DNA virus MAGs from the Global Ocean Eukaryotic Viral database^45^, which includes *Nucleocytoviricota* genomes and 111 genomes belonging to the recently-discovered *Mirusviricota* lineage. Additionally, we randomly selected 250 complete genomes of large *Caudovirales* (jumbo bacteriophages) from the INPHARED database^46^ (5Jan2023 version). We included these groups of viruses for training because they commonly encode genes with matches to the GVOG profiles^40^ and therefore likely to be falsely classified as giant viruses.

To avoid the inclusion of identical or highly similar genomes, we performed genome dereplication using dRep v3.2.2^47^ (dereplicate command, with parameters -l 5000 -- ignoreGenomeQuality -pa 0.95 --SkipSecondary) . We arrived at a nonredundant set of 1,912 viral genomes (1,551 *Nucleoviricota*, 111 *Mirusviricota*, and 250 jumbo phage sequences) for downstream training and testing.

### Training set and independent test set

We split the compiled database into two independent genome sets for the purposes of training and benchmarking. We randomly assigned 80% of the genomes in each viral group (each family of the *Nucleoviricota*, *Mirusviricota*, and jumbo phages), totalling 1,531 sequences, to the training set. The remaining 20% of the genomes (381 in total) were assigned to the test set. The test set contained sequences of the from the *Mirusviricota*, jumbo phages, and 10 virophages to serve as non-giant virus genomes. We previously delineated taxonomic classification for the *Nucleocytoviricota*^7^, and here we used the same nomenclature in training.

Giant virus genomes assembled from metagenomes can be fragmented and incomplete. To simulate these incomplete cases, we utilized a custom Python script that generated fragmented genomes at random completeness levels (genome_fragmentizer.py at https://zenodo.org/records/10085666). For each of the initial giant virus genomes (n = 1,242), we generated 2 fragmented sequences at random completeness levels (compared to the initial genome) ranging from 23 to 99%. For each of the *Mirusviricota* and jumbo phages genomes (n = 289), we generated 1 fragmented version at random completeness level, ranging from 29 to 99%. Due to the limited number of *Pokkesviricetes* incertae sedis genomes available (only 2 in our dataset), we created 6 fragmented versions for each of them to introduce variability. Collectively, this process resulted in a total of 4,316 sequences at varied completeness levels for the training dataset. The detailed taxonomic classification and completeness level of each genome sequence are detailed in Table S1.

We implemented a similar fragmentation process for the testing set. For each of the initial 381 genomes reserved for benchmarking, we generated 1 fragmented version at random completeness level. Additionally, for each of the 9 reference genomes isolated from culture and assembled into a single contig, we generated 6 fragmented versions to introduce more variability. This resulted in a total of 823 sequences for benchmarking, with completeness values spanning from 33% to 100%. Table S2 detailed the taxonomic assignments and completeness levels of the sequences in the testing set.

### Giant Virus Orthologous Groups (GVOGs) as Features for Classification Models

Hidden Markov Models (HMMs) of 8,863 protein families found in giant virus genomes, which we refer to as giant virus orthologous groups (GVOGs) were downloaded from the GVDB. Details regarding GVOG construction have been previously described^11^. Given that there is a limited number of genes shared across different giant virus orders, we screened for GVOGs that are found in at least 25% of the genomes within each order. We arrived at a set of 625 GVOGs that were broadly represented across different orders of the Nucleocytoviricota that we used for the first round of model training.

### Processing sequences for training

To prepare training data for model construction, we first predicted proteins from genomes using Prodigal^48^ V2.6.3, with default parameters. Next, we compared predicted proteins to the pre-specified set of GVOG HMMs using the hmmsearch command in HMMER3 3.3^49^, with an e-value threshold of 1e-10. Additionally, we calculated the GC content of each genome sequence through a custom Python module. Although genome size can be viewed as a distinguishing feature of some giant virus lineages, we excluded this feature because we sought to develop an approach that could be used for incomplete genomes. These steps collectively generated a feature matrix to be passed to the RF classifiers.

### Construction of classification models

The RF algorithm was applied using the sci-kit learn library ^50^ v1.2.1 in Python v3.8.18. Training was performed separately for the Order and Family levels. In the first round of training, all 625 prevalent GVOGs (present in at least 25% of genomes in each giant virus order) and GC content of the sequences were used as features. We first perform randomized search cross validation using Scikit-Learn’s RandomizedSearchCV method. This involved defining a grid of hyperparameters across a broad range, randomly sampling values from this grid, and assessing the performance of the models for each combination of values. Based on the best hyperparameter values provided by random search, we defined a new hyperparameter grid and and selected optimal hyperparameters for classification models through 10-fold grid search cross-validation (GridSearchCV).

To evaluate how the models’ accuracy varied with different training test size, we split the data set into training and cross-validation folds through 10-fold cross-validation. Subsets of sequences ranging from 25 to 100% of the training set size were drawn from each training fold, and a model was trained through grid search cross-validation on each subset. The mean and 95% confidence interval for training and cross-validation accuracies across all folds at each number of sequences were reported. The accuracy metric, an evaluation measure of model performance, was calculated as follows:

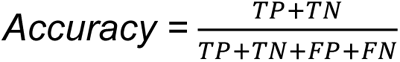

where: TP = True positive; FP = False positive; TN = True negative; FN = False negative.

### Reducing the number of GVOG features

We first investigated whether the number of GVOG features used in training significantly influenced prediction accuracy using Recursive Feature Elimination (RFE) with cross-validation. RFE fits the model multiple times and removes the weakest features until it reaches a given number of features. Best subsets of features at different sizes were scored and reported. We examined the collinearity between GVOG features by performing hierarchical clustering on the Spearman rank-order correlations using the spearmanr() function from scipy.stats.

To identify features that were more important to the performance of classification models and reduce the number of features required for classification, we calculated feature importance using the RF’s fitted attribute *feature_importances_*. This measures the importance of a feature by computing the mean and standard deviation of accumulation of the impurity decrease within each tree when including that feature. In addition, we performed an alternative method, permutation feature importance, to inspect the model. Permutation feature importance measures the decrease in a model’s performance score when a single feature value is randomly shuffled. We calculated the permutation importances on a held-out set to determine which features most significantly contribute to the model’s generalization capabilities. It is worth noting that the GVOG data exhibited collinearity (Fig. S2). When features are highly correlated, permuting a single feature may not significantly affect the model’s performance as the model can access the same information through its correlated feature. This can reduce the importance value of these features, even though they may actually be important. To address this, we employed hierarchical clustering with Ward’s linkage to group features and retained one feature from each cluster. Subsequently, we calculated the permutation importance of the selected set after removing redundant features.

After identifying the set of most important features using each method, we subsequently retrain RF models using new feature matrices. We performed hyperparameter tuning using random search and grid search as described above. The performance of the two sets of models at the order and family level was estimated using 10-fold nested cross-validation. In this procedure, we selected models through grid search cross-validation within an outer cross-validation loop. For each iteration of the outer loop, we constructed and selected the best model using GridSearchCV, and then evaluated this model on the test set of the outer fold. Nested cross-validation estimates how effectively a model trained with a specific strategy will generalize to previously unseen data. The final set of classification models were selected based on performance.

### Evaluating classification model performance with an independent test set

The classification models were tested against an independent test set, excluded from model generation, of 823 sequences at varied genomic completeness, ranging from 33% to 100%. The predicted taxonomic classification was compared to the actual classification for these sequences to estimate accuracy and generate confusion matrices and classification reports. In the classification reports, in addition to accuracy, three other metrics were used to evaluate the performance of the classification models: precision (correctness), recall, and F1 score. F1 score is a widely used metric to evaluate multiclass classification problems as it balances precision and recall. The metrics were calculated as follows:

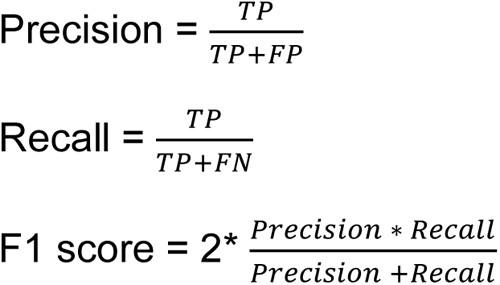

where: TP = True positive; FP = False positive; TN = True negative; FN = False negative.

### Average amino acid identity (AAI) calculation

It can also be useful to know the best match of a query viral genome against a reference database, and so we also implemented a one-way average amino acid identity (AAI) search in TIGTOG. AAI between the input genomes and a custom reference database can be requested using the -a flag. Rather that include the entire set of giant virus genomes available in the GVDB, we included only one representative from each genus classified in this database, which was chosen based on N50 contig length^11^. The module employs one-way LAST searches^51^ (parameter -m 500) of predicted proteins in input sequences against our database and calculates the AAI and alignment fraction (AF) between all genome pairs. To avoid partial matches, input genomes having an AF < 20 were considered to have an AAI of 0.

## Data and Code Availability

The custom script for genome fragmentation and the genome sets used for model training and testing are available on Zenodo at https://zenodo.org/records/10085666. Source code and instructions for TIGTOG are available on Github at https://github.com/anhd-ha/TIGTOG.

## Author information

### Contributions

FOA conceived the project. ADH and FOA designed algorithms and databases. ADH performed analysis and validation and developed the software. ADH and FOA wrote the manuscript.

## Acknowledgements

We would like to thank Carolina Martinez Gutierrez for assistance with genome fragmentation. We thank members of the Aylward Lab for helpful comments. This work was performed using compute nodes available at the Virginia Tech Advanced Research and Computing Center. This work was supported by grants from the National Science Foundation (CAREER-2141862 to F.O.A.) and the National Institutes of Health (1R35GM147290-01 to F.O.A).

