## Supplementary Figures for "Automated classification of giant virus genomes using a random forest model built on trademark protein families"

**Figure S1. Distinct protein family profiles in *Nucleocytoviricota* families.** The y-axis denotes the family assignments of giant virus representative genomes, color-coded by family. The x-axis shows different GVOGs. The Not_GV group includes *Mirusviricota* and jumbo phage genomes.

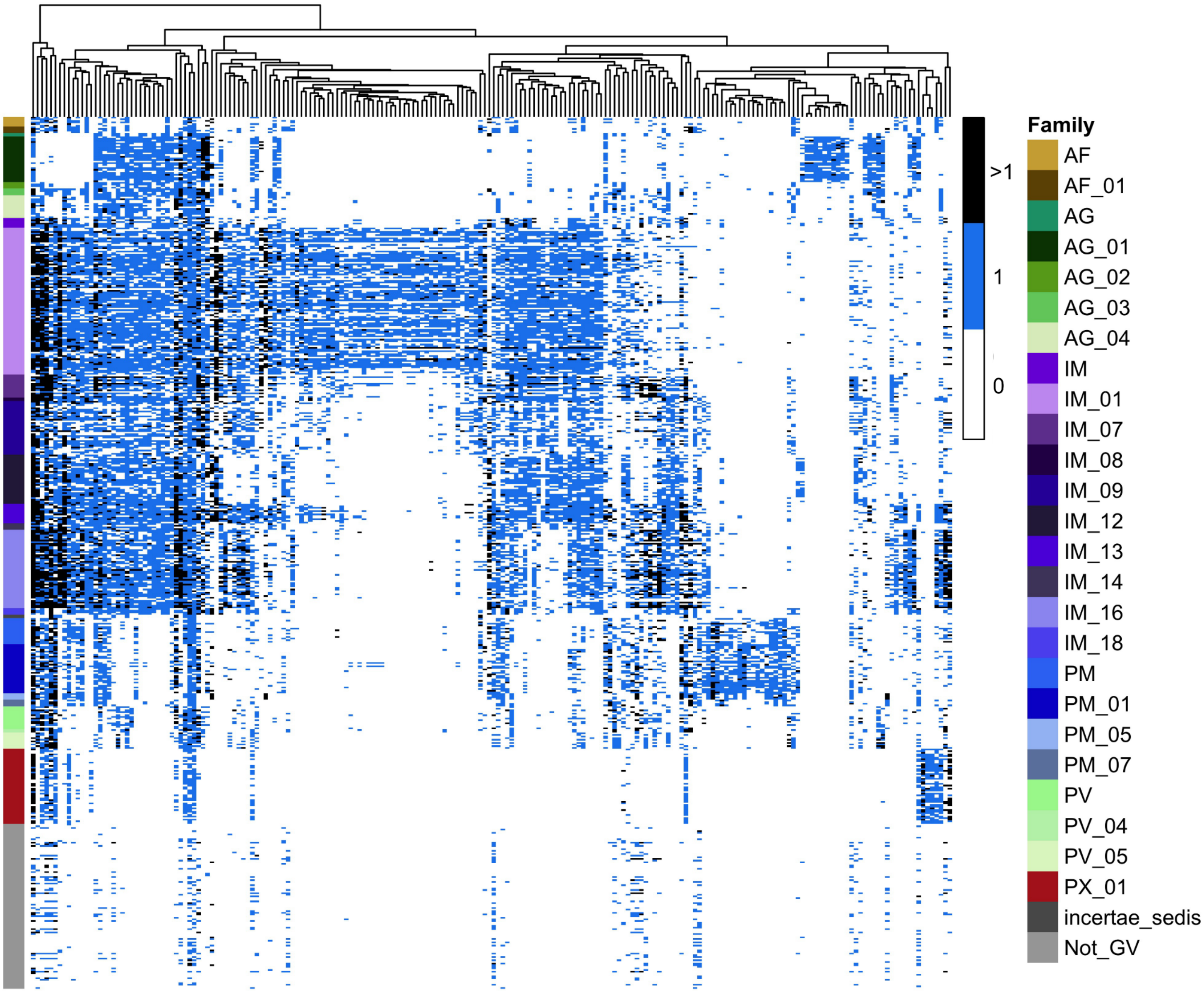

**Figure S2. Heatmap of correlated features in the initial 625 GVOG feature set.** Hierarchical clustering were performed on the Spearman rank-order correlations using Ward's linkage.

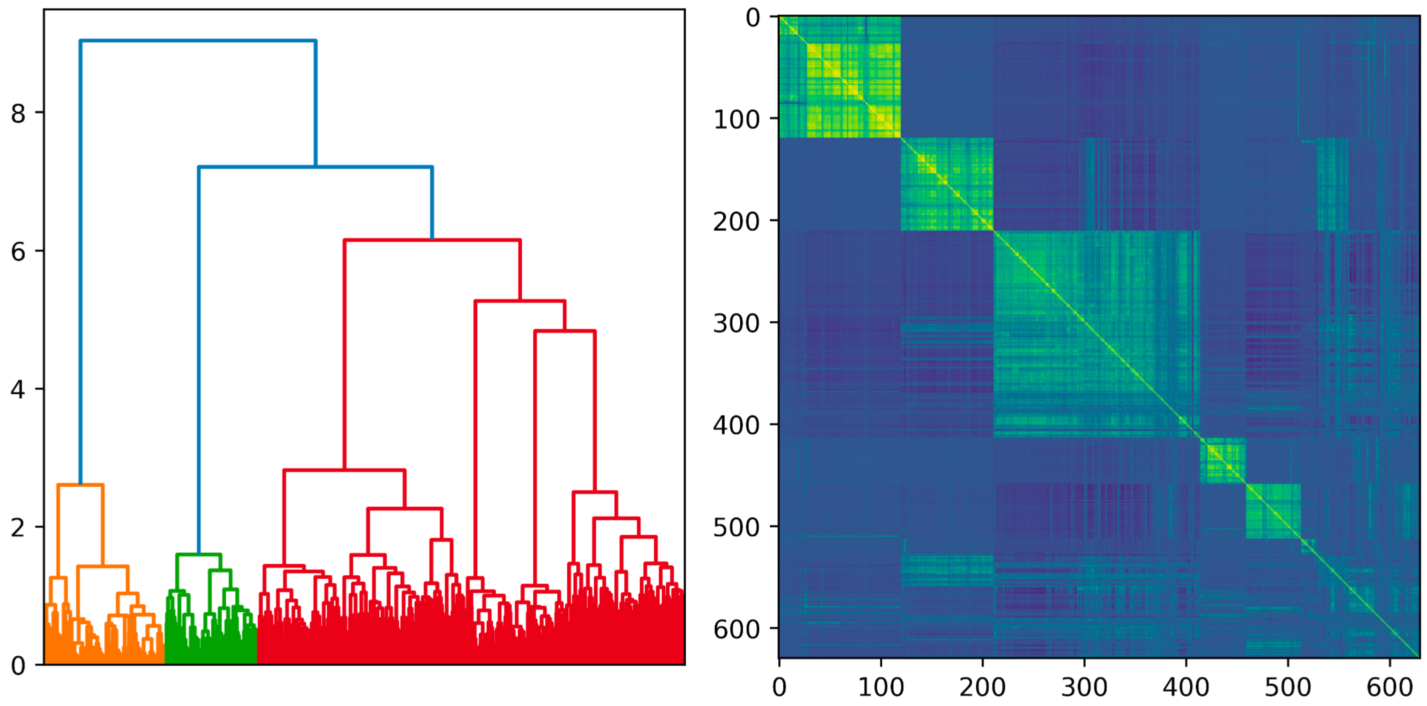

**Figure S3. MDI-based feature importances for the 15 most important features in the classification model at the order level (top) and family level (bottom).**

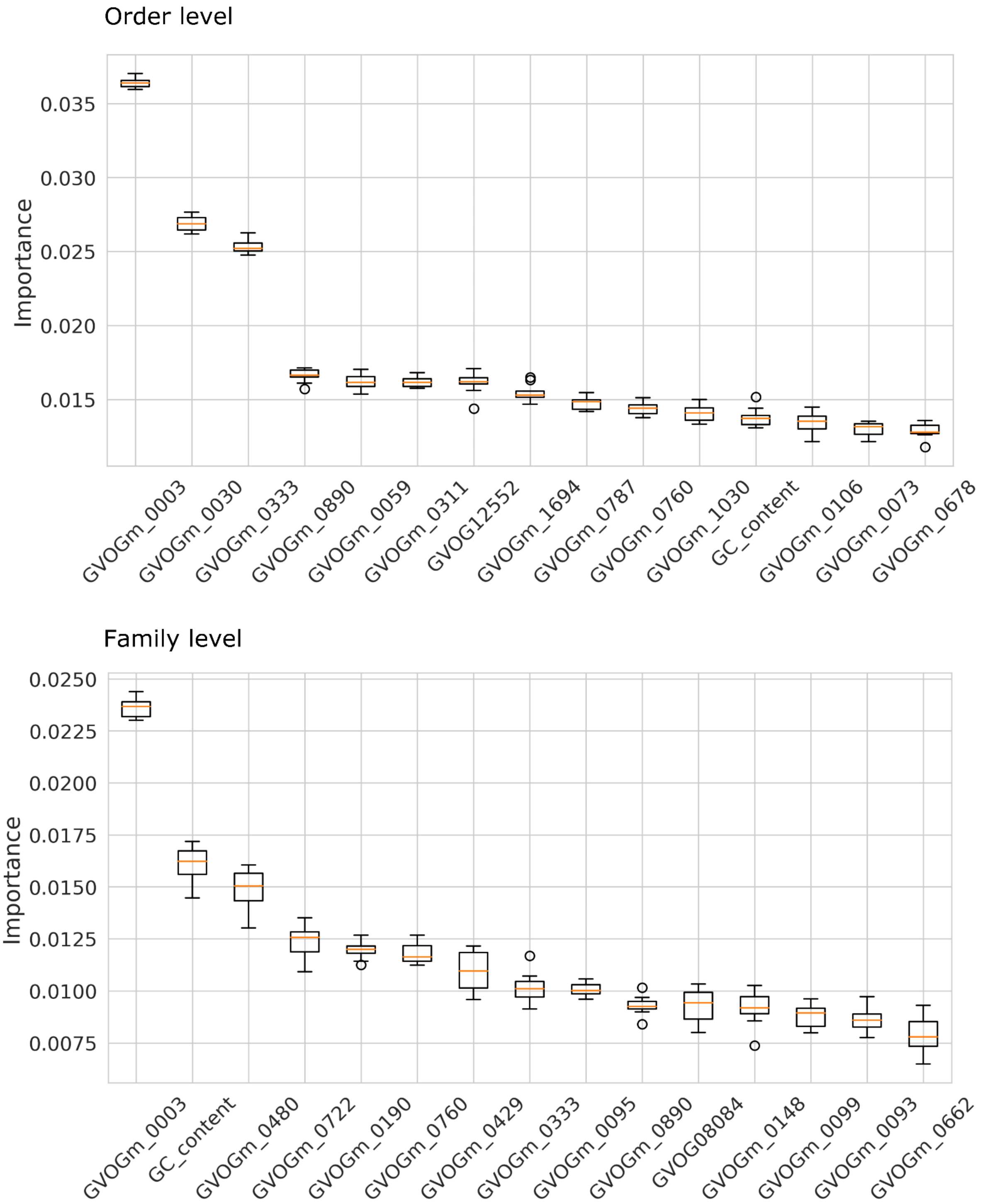

**Figure S4. Performance of models feature sets selected by two feature importance mechanisms.** Model performance was estimated using 10-fold nested cross-validation.

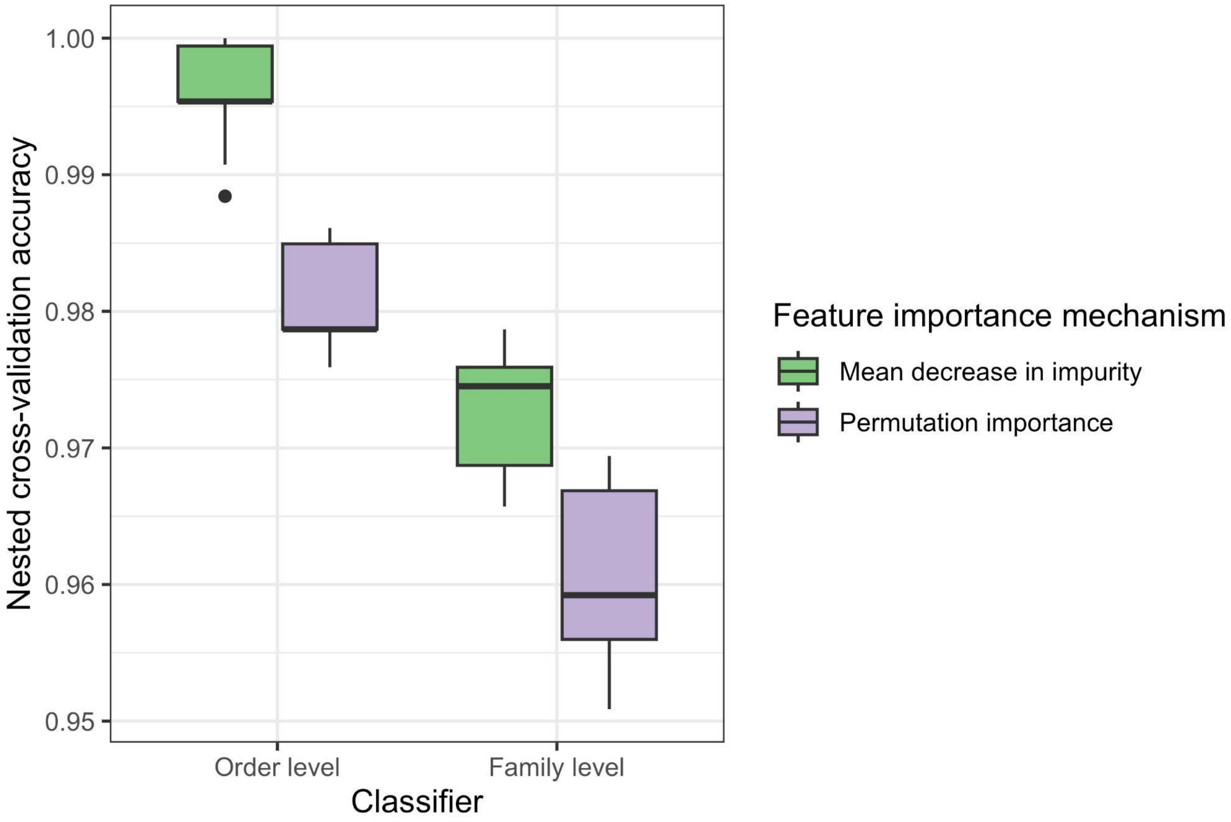

**Supplemental Data S1.** GVOG features used for models at the order and family levels. Features were selected using RF's MDI-based importance.

**Supplemental Data S2.** Summary of viral genomes included in model training and independent testing.

**Supplemental Data S3.** Classification report at the family level.

**Supplemental Data S4.** Summary of giant virus genomes included in the custom AAI database.
